# OrganelX Web Server for Sub-Peroxisomal and Sub-Mitochondrial protein localisation

**DOI:** 10.1101/2022.06.21.497045

**Authors:** Marco Anteghini, Asmaa Haja, Vitor AP Martins dos Santos, Lambert Schomaker, Edoardo Saccenti

## Abstract

Computational approaches for sub-organelle protein localisation and identification are often neglected while general methods, not suitable for specific use cases, are promoted instead. In particular, organelle-specific research lacks user-friendly and easily accessible computational tools that allow researchers to perform computational analysis before starting time-consuming and expensive wet-lab experiments. We present the Organelx e-Science Web Server which hosts three sequence localisation predictive algorithms: In-Pero and In-Mito for classifying sub-peroxisomal and sub-mitochondrial protein localisations given their FASTA sequences, as well as the Is-PTS1 algorithm for detecting and validating potential peroxisomal proteins carrying a PTS1 signal. These tools can be used for a fast and accurate screening while looking for new peroxisomal and mitochondrial proteins. To our knowledge, this is the only service that provides these functionalities and can fasten the daily research of the peroxisomal science community.

## 1 Introduction

Signatures in the amino acid composition of proteins have been associated with protein domains, families functional sites and their subcellular localisation [1–4]. This information can be combined with machine learning (ML) approaches to develop prediction tools, that nowadays are easily findable and accessible [5–8].

Moreover, ML and deep-learning (DL) approaches are often used to encode protein sequences, thus showing promising results in several tasks, including subcellular classification [9–15]. Unified Representation (UniRep) [9] and the Sequence-to-Vector (SeqVec) [10] are two of most promising and already used DL-embeddings embeddings.

UniRep [9] provides amino-acid embedding containing meaningful physicochemically and phylogenetic clusters and proved to be efficient for distinguishing proteins from various SCOP (structural classifications of proteins) classes. SeqVec showed similar results and optimal performance for predicting subcellular localisation [10]. However, their potential has been explored for highly specific tasks, such as sub-organelle localisation, just recently [16]. In particular, their usage can be adapted for sub-peroxisomal and sub-mitochondrial protein localisation.

Peroxisomes and mitochondria are ubiquitous organelles surrounded by a single biomembrane (peroxisomes) or a double biomembrane (mitochondria) that are relevant to many metabolic and non-metabolic pathways [17, 18]. However, the full extent of the functions of peroxisomes, mitochondria and their shared pathways is still largely unknown [19] and the discovery of new peroxisomal and mitochondrial proteins can facilitate further knowledge acquisition.

The OrganelX Web Server, available at https://organelx.hpc.rug.nl/fasta/, hosts two algorithms designed to classify sub-peroxisomal (In-Pero) and sub-mitochondrial (In-Mito) proteins localisations given their FASTA sequences. In addition, it is possible to perform a specific protein targeting signal (PTS) research in the sequence (Is-PTS1). Our tool consents to detect and validate on the PTS1 associated with peroxisomal proteins. The PTS1 is present in the C-terminal part of the protein sequence. It is defined as the final dodecamer with a focus on the terminal tripeptide [20]. An overview of the functionalities available in the Web Server is shown in Figure 1.

**Figure 1.**
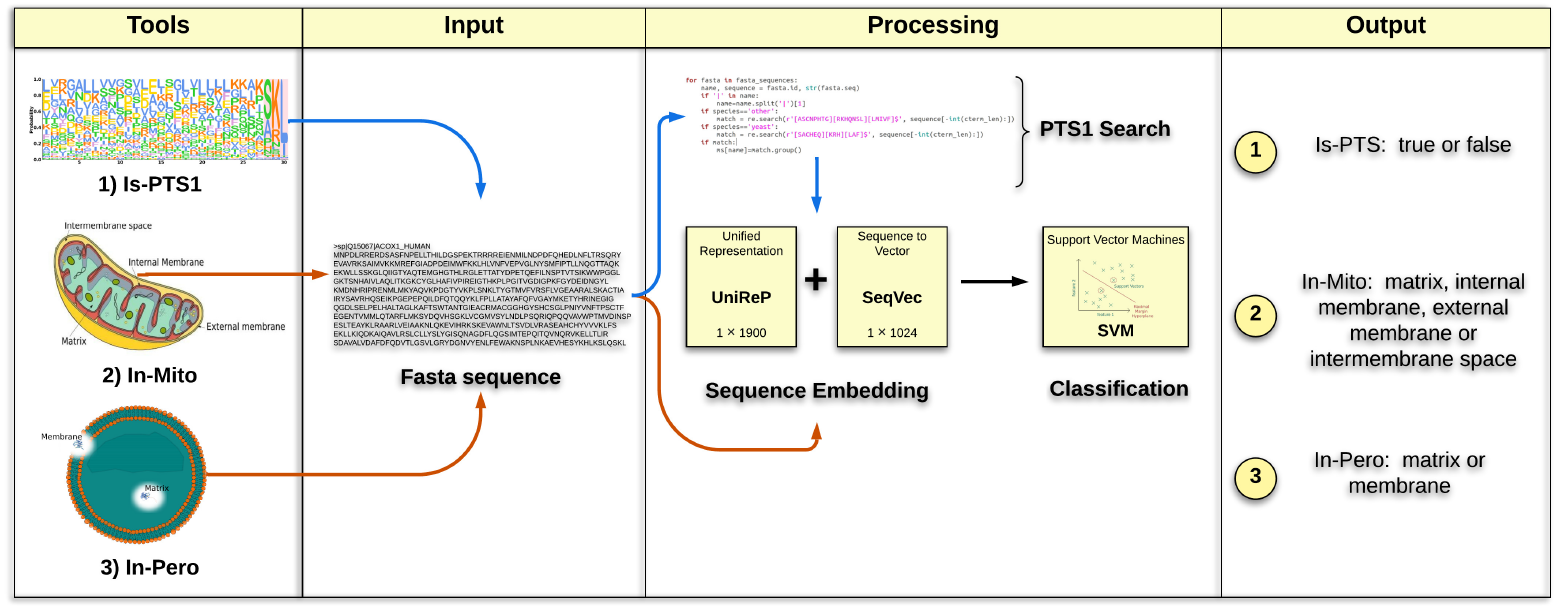
Web Server tools overview. The 4 columns show the typical workflow which consists of: (1) tool selection (Is-PTS1, In-Pero, In-Mito); (2) input a FASTA file containing one or more sequences; (3) processing via sequence embedding representation and SVM classification; (4) output generation.

To our knowledge, there is no service that automatically performs In-Pero and In-Mito classification as well as Is-PTS1 detection. These predictions are noticeably useful for peroxisomal-related research [21].

## 2 OrganelX functionalities and examples

### Input

The input is a FASTA text file containing one or more protein sequences. Each sequence begins with a single-line description, followed by sequence data. The single-line description consists of **>sp**|**ID**|**Desc** symbols, where **>sp** is a fixed prefix, **ID** is the sequence name, and **Desc** is a descriptive text, followed by tokens of the FASTA sequence on the next lines.

### Output

The output file is a plain text file containing the classification results. The results of each class are represented by the predicted class name followed by ‘:’ and the IDs of the corresponding classified entries.

### Processing Methods

The three classification algorithms start representing the sequence as a concatenation of two deep learning-based sequence embeddings, namely UniRep and Seqvec [9, 10]. The classification is performed by Support Vector Machines (SVM), where the In-Pero and In-Mito algorithms classify among sub-organelle classes. There are two classes (matrix and membrane) for peroxisomal proteins and four classes (matrix, inner-membrane, intermembrane, outer-membrane) for mitochondrial proteins. The third algorithm detects the PTS1 signal by matching a regular expression. Adopting a similar workflow that starts with the embedding representation and performs an SVM based classification, the Is-PTS1 validation algorithm classifies true and false PTS1.

### Validation

The algorithm was validated on two validation sets. Data were downloaded from Uniprot [22] on the 25/02/2021. For validating In-Pero, we queried for reviewed proteins with a clear sub-peroxisomal annotation in membrane (“SL-0203” and “GO:0005778”) or matrix (“SL-0202” and “GO:0005782”). We then clustered the query results for 40% of sequence identity with CD-hit [23] and removed the sequences overlapping with our training set, obtaining 85 membrane proteins and 59 matrix proteins. Our model obtained an accuracy of 0.88. We also tested the Is-PTS1 predictor retrieving in a similar way 15 true and 15 false, where true refers to a peroxisomal protein carrying the PTS1 signal and false refers to a peroxisomal protein without PTS1 signal. The performance in terms of accuracy is 0.83. The in-depth analysis of the In-Mito predictor is available in the original paper [16]. Datasets are available at https://github.com/MarcoAnteghini/.

### Examples

In order to submit a job, the user must specify an arbitrary username, his email address (optional) and upload a FASTA file. The computation time changes depending on the file size and the traffic on the website. The user can either wait for the results via email, or simply refreshing the result web page. An example is visible at https://organelx.hpc.rug.nl/fasta/test_example, where also a FASTA file can be downloaded.

## Acknowledgements

We thank the Center for Information Technology of the University of Groningen for providing access to the Peregrine high performance computing cluster. The e-Science server was realized with the support and nurturing of Ger Strikwerda.

## Author Contribution

- Category 1

Conception and design of study: MA, AH, ES; acquisition of data MA, AH; analysis and/or interpretation of data: MA, AH, ES, VMS, LS.

- Category 2

Drafting the manuscript: MA, AH; revising the manuscript critically for important intellectual content: LS, ES, VMS.

- Category 3

Approval of the version of the manuscript to be published: MA, AH, VMS, LS, ES.

